# GGAssembler: precise and economical design and synthesis of combinatorial mutation libraries

**DOI:** 10.1101/2023.05.18.541394

**Authors:** Shlomo Yakir Hoch, Ravit Netzer, Jonathan Yaacov Weinstein, Lucas Krauss, Karen Hakeny, Sarel Jacob Fleishman

## Abstract

Golden Gate assembly (GGA) can seamlessly generate full-length genes from DNA fragments. In principle, GGA could be used to design combinatorial mutation libraries for protein engineering, but creating accurate, complex, and cost-effective libraries has been challenging. We present GGAssembler, a graph-theoretical method for economical design of DNA fragments that assemble a combinatorial library that encodes any desired diversity. We used GGAssembler for one-pot *in vitro* assembly of camelid antibody libraries comprising >10^5^ variants with DNA costs <0.007$ per variant and dropping significantly with increased library complexity. >93% of the desired variants were present in the assembly product and >99% were represented within the expected order of magnitude as verified by deep sequencing. The GGAssembler workflow is, therefore, an accurate approach for generating complex variant libraries that may drastically reduce costs and accelerate discovery and optimization of antibodies, enzymes and other proteins. The workflow is accessible through a web interface at https://github.com/Fleishman-Lab/GGAssembler/blob/master/example/colab_oligos_design.ipynb.

## Introduction

Over the past decade, high-throughput screening of protein variant libraries and deep sequencing have revolutionized protein characterization, engineering, and design(Wójcik et al. 2015; Markel et al. 2020). Common library-synthesis approaches introduce random or systematic single-point mutations(Alejaldre, Pelletier, and Quaglia 2021). However, significant gains in protein activity and stability often require multiple simultaneous mutations(Khersonsky et al. 2018; Whitehead et al. 2012), calling for gene-synthesis approaches that introduce combinatorial mutations. Such multipoint combinatorial libraries can be designed based on previous experimental screens(Whitehead, Baker, and Fleishman 2013; Whitehead et al. 2012; Fowler et al. 2010) and increasingly based on computational design calculations(Guntas, Purbeck, and Kuhlman 2010; Weinstein et al. 2023; Treynor et al. 2007; Listov et al. 2024). However, synthesizing combinatorial libraries accurately and economically is challenging due to the large size of a typical binding or enzyme active site and the distribution of active-site positions across multiple noncontiguous epitopes. An illustration of the potential complexity of effective variant libraries is provided by our recent computational active-site library design studies from which thousands of diverse and functional variants were isolated in a single experiment(Lipsh-Sokolik et al. 2023; Weinstein et al. 2023). In these studies, atomistic design calculations defined the desired diversity across dozens of positions, including, in one case, through combinatorial insertions and deletions at an enzyme active site(Lipsh-Sokolik et al. 2023). To synthesize such complex yet completely defined variant libraries, we considered different cloning strategies that would be general and economical, enable a high level of control over the location and identity of mutations and the ratio of mutations to the wild-type parental identities, and exhibit low representation bias among the variants.

Given these demands, we focused on Golden Gate Assembly (GGA)(Engler, Kandzia, and Marillonnet 2008; Engler et al. 2009). GGA employs Type IIs restriction enzymes, such as *BsaI*, that cleave outside their DNA recognition sequence. Following processing by the Type IIs restriction enzyme, the recognition site is eliminated, leaving terminal overhangs that can be programmed to encode immutable (non diversified) amino acid positions. Subsequent ligation reactions of the processed fragments then seamlessly combine them into full-length genes(Engler, Kandzia, and Marillonnet 2008; Engler et al. 2009). In principle, GGA enables scarless gene construction of any size from small fragments(Andreou and Nakayama 2018; Püllmann et al. 2019; Sarrion-Perdigones et al. 2011, 2013; Vazquez-Vilar et al. 2017; Weber et al. 2011). High-fidelity GGA, however, is governed by several factors that limit the number and identity of overhangs: the nonspecific (star) activity of the restriction enzyme(Mayer 1978), the sensitivity of the DNA ligase to mismatches(Lohman et al. 2016), and the relative propensity of DNA overhangs to anneal under different reaction conditions. These limitations forced previous GGA applications to use iterative DNA assembly methods with a restricted set of overhangs(Andreou and Nakayama 2018; Püllmann et al. 2019; Sarrion-Perdigones et al. 2011, 2013; Vazquez-Vilar et al. 2017; Weber et al. 2011). As a step to address these limitations, recent studies have examined the results of GGA under different experimental conditions and expanded the set of allowed overhangs(Potapov, Ong, Kucera, et al. 2018; Pryor et al. 2020; Potapov, Ong, Langhorst, et al. 2018). In principle, these developments allow the assembly of dozens of fragments for combinatorial gene synthesis(Potapov, Ong, Kucera, et al. 2018; Pryor et al. 2020). Despite much progress(Püllmann et al. 2019; Pryor et al. 2020; Kirby et al. 2021; Daffern et al. 2023), however, methods for economical and accurate construction of variant libraries from dozens of fragments have not yet been demonstrated.

We present a new method, called GGAssembler, for the economical design of DNA fragments that assemble into a gene library which encodes any specified diversity with minimal representation bias. We recently used GGAssembler to synthesize two complex libraries both encoding millions of combinatorial mutants at 14 positions in the GFP chromophore-binding pocket with a total DNA cost of $455, translating to 4.2 * 10^−3^ ₵ per variant(Weinstein et al. 2023). We now demonstrate that this approach can accurately, efficiently, and economically encode complex multipoint mutation libraries of camelid single-domain (vHH) antibodies comprising hundreds of thousands of variants. GGAssembler can be applied, in principle, to genes of all sizes, but the relatively small size of the vHH gene allows us to rigorously demonstrate completeness, high accuracy, and low representation bias in the final assembly product. GGAssembler is freely available at https://github.com/Fleishman-Lab/GGAssembler and can be accessed through an online notebook at https://github.com/Fleishman-Lab/GGAssembler/blob/master/example/colab_oligos_design.ipynb.

## Results

### A general method to design combinatorial diversity

We developed GGAssembler to provide an economical approach to combinatorial library assembly while allowing nearly complete freedom to decide on mutation types, positions, and the size of the library. Critically, GGAssembler guarantees a lower bound on the fidelity of the GGA reaction. To estimate the probability that a given set of overhangs assembles with high fidelity and accuracy, we leverage empirically derived overhang ligation fidelity (**Eq. 1**) and efficiency (**Eq. 2**) estimates(Potapov, Ong, Kucera, et al. 2018; Pryor et al. 2020; Potapov, Ong, Langhorst, et al. 2018). The cost associated with each DNA fragment is the number of nucleotides required to produce all the diversity encoded in that fragment. For example, the cost associated with a 20nt DNA fragment that encodes a single mutated amino acid position that requires two codons would be 40nt (the length of the fragment times the encoded variability). Adding an extra codon at the mutated position would increase the cost to 60nt, whereas adding another mutated position would multiply the cost by the number of codons in that position. GGAssembler uses degenerate codons to encode as much desired diversity as possible at a minimal DNA cost while completely excluding undesired amino acid identities and stop codons.

The GGAssembler workflow comprises three steps as schematically represented in **Figures 1 and 2** and **Supplementary Figure 1**. We start by defining the amino acids that encode the desired diversity with an optional codon-compression step that uses degenerate codons(Knuth 2000) (Methods). This step excludes all unspecified amino acids and stop codons. In the next step, we employ a graph-theoretical cost-optimization algorithm to compute alternative solutions that vary in cost and number of overhangs. To accomplish this, we build a graph in which nodes represent restriction enzyme cleavage sites, and edges connect all pairs of sites that a single DNA oligo can traverse. To construct this graph, we start with a user-defined DNA sequence of the target gene prior to amino acid diversification, a list of desired amino acid mutations, the Type IIs restriction enzyme, the minimum required level of ligation efficiency, and criteria that define acceptable DNA fragments, such as minimal and maximal oligo lengths. We then search for potential cleavage sites given the size of the overhang generated by the restriction enzyme. To serve as a cleavage site, the DNA region encompassing the nucleotide and the overhang generated by the restriction enzyme must be immutable (it cannot encode diversity), and the overhang must exhibit at least the desired ligation efficiency according to empirical measurements(Potapov, Ong, Kucera, et al. 2018; Pryor et al. 2020; Potapov, Ong, Langhorst, et al. 2018). We add sites that fulfill these requirements as nodes to the graph with their associated overhangs. An edge connects each pair of nodes if the two overhangs do not ligate to one another. We assign each edge a weight based on the number of nucleotides (proportional to the cost) required to generate the variability in the DNA fragment it represents.

**Figure 1.**
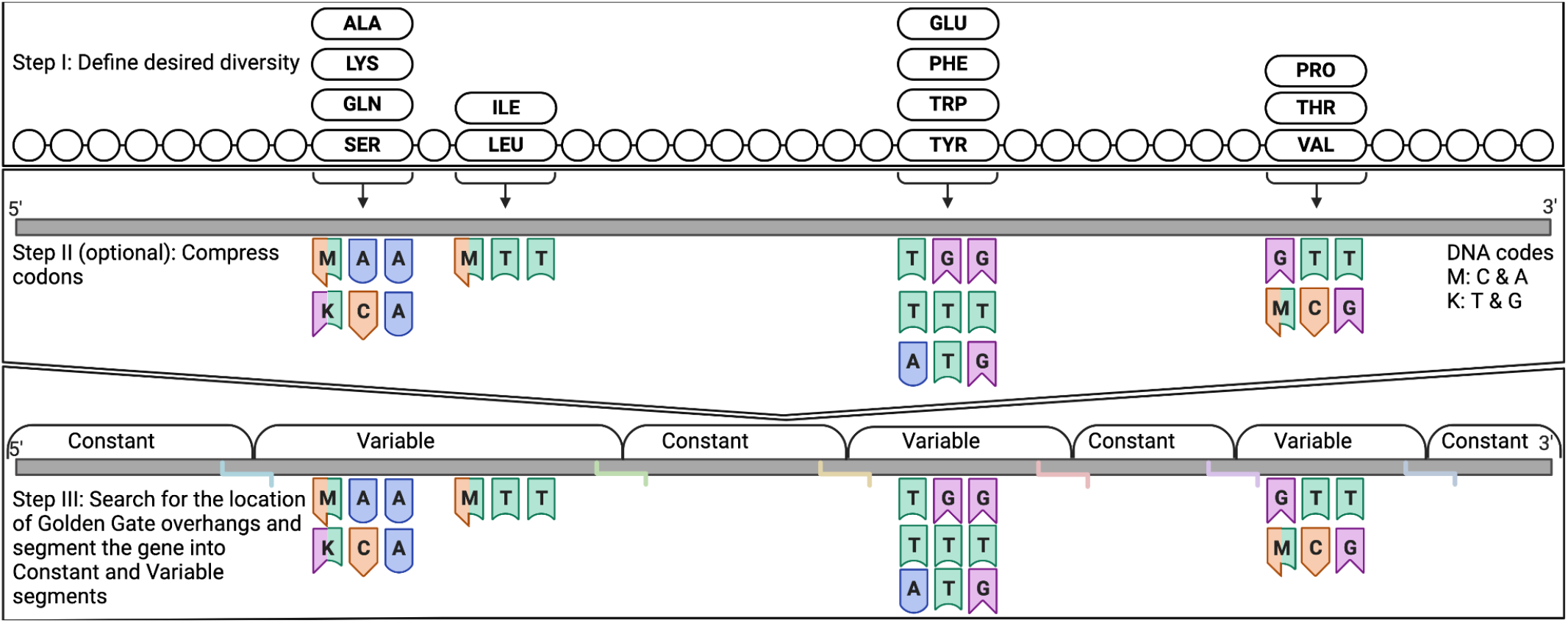
Key steps in the GGAssembler computational workflow. Step I: Provide wild type DNA sequence and the required diversity in each position. Optional step II: Apply codon compression for the required diversity in each position. Step III: Apply a shortest-path and rainbow shortest-path search to find an economical segmentation of the DNA that maintains GGA reaction fidelity above a given threshold.

**Figure 2.**
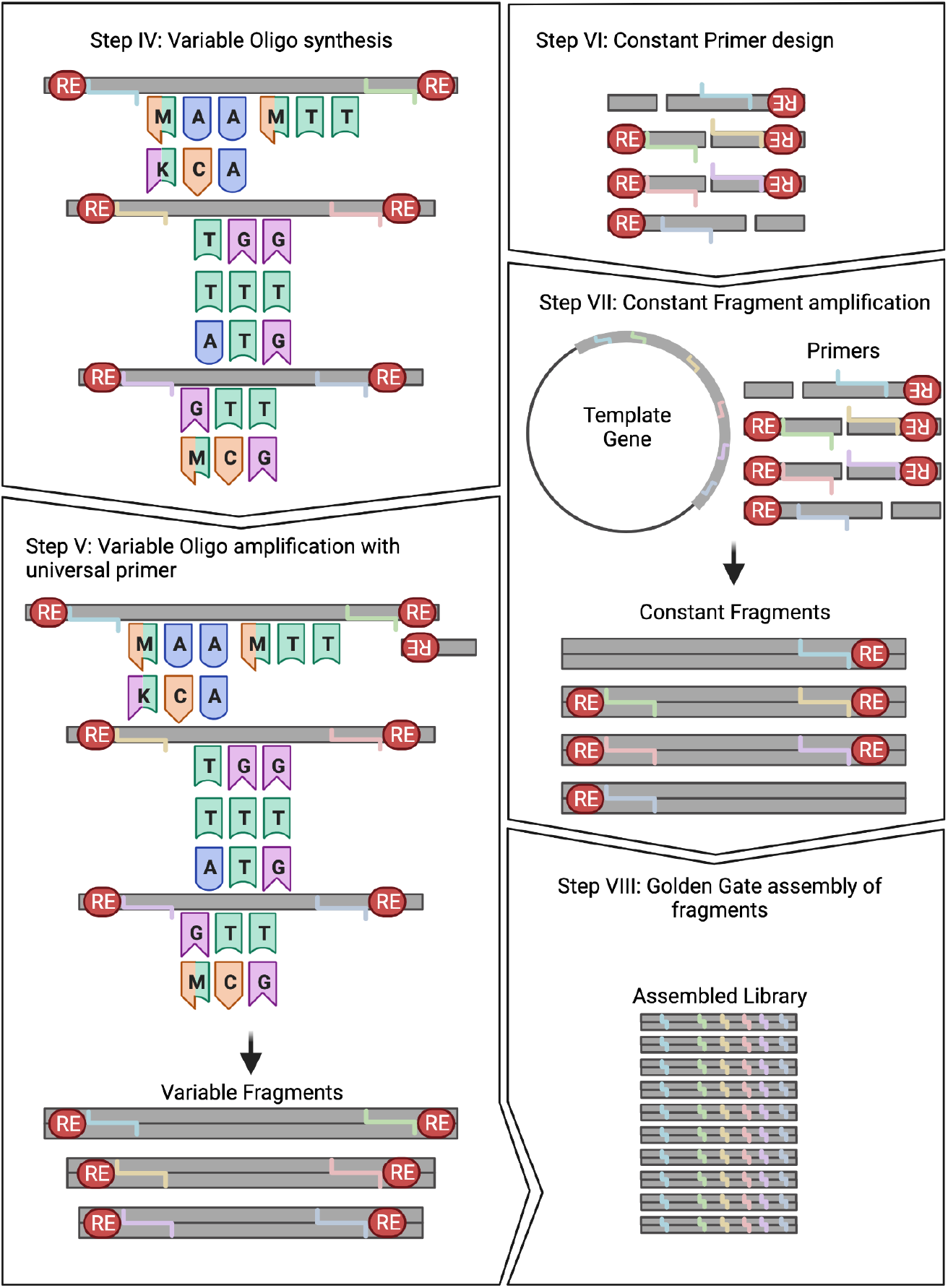
Key steps in the GGAssembler experimental workflow. Step IV: Order thevariable segments as ssDNA. Step V: Amplify variable segments ssDNA to construct dsDNA. Step VI: Manually design and order primers for constant segments based on the output EMBL sequence file. Step VII: Amplify and purify constant segments from the input gene template. Step VIII: Execute GGA following one of several established protocols(Potapov, Ong, Langhorst, et al. 2018; Pryor et al. 2020, 2022; Sikkema et al. 2023).

A critical confounding factor in combinatorial assembly of multiple fragments that is not addressed by the scheme above is that non-adjacent overhangs might inadvertently exhibit high ligation efficiency resulting in undesired assembly products (see **Figure 3** for examples). Some misassembled products may be shorter than the desired one and would therefore be preferentially amplified in downstream PCRs. Furthermore, some cases may result in loops of misassembled concatemers. To ensure strict assembly only of the desired product, we must ensure fidelity above the user-defined threshold. Providing the most economical solution while maintaining fidelity above the provided threshold is referred to as the constrained shortest-path problem, which is a recognized NP-hard problem (Lozano and Medaglia 2013). To overcome the hardness barrier we employ randomized color-coding(Alon, Yuster, and Zwick 1995) in which nodes that represent overhangs that are predicted to ligate at high efficiency are colored identically. We then apply a vertex rainbow shortest-path algorithm that ensures that valid paths include each color at most once, resulting in vertex rainbow paths(**Figure 3**). Selecting rainbow shortest-path solutions guarantees the assembly of an economical solution that excludes undesired products.

**Figure 3.**
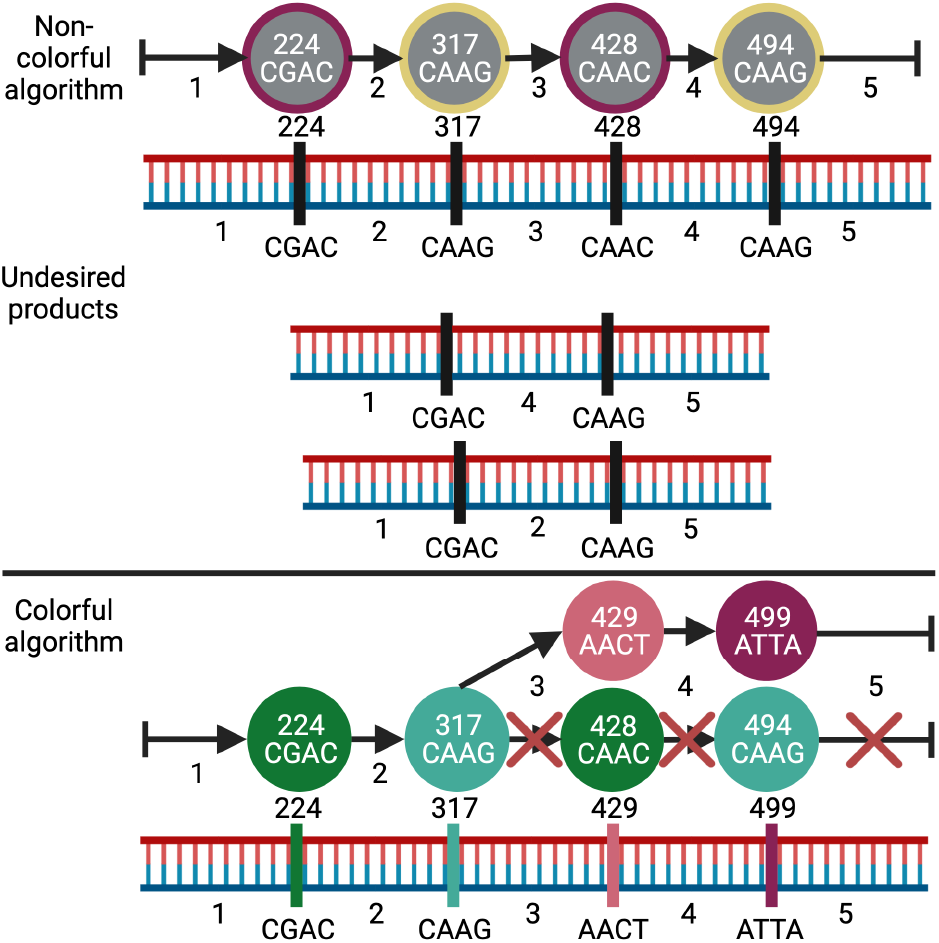
Graph coloring in GGAssembler eliminates undesired assemblies. A subset of the path common to all shortest paths in a previously designed GFP library that encodes >10^7^ active-site variants(Weinstein et al. 2023). (top) Nodes marked by cleavage-site location and overhang are colored identically if the overhangs ligate. Two nodes with the same overhang (CAAG, positions 317 and 494) are marked yellow, and nodes that are not Watson-Crick pairs but are predicted to exhibit high-efficiency ligation(Potapov, Ong, Kucera, et al. 2018; Pryor et al. 2020; Potapov, Ong, Langhorst, et al. 2018) are in purple. All shortest-path solutions would result in misassembled products. In this case, segments 1 and 4 are complementary as are 2 and 5, leading to specific undesired products shown in the figure. In addition, the end of segment 3 can ligate to the beginning of segment 2 leading to an infinite number of different undesired products (not shown). (bottom) A rainbow shortest-path search excludes undesired solutions, forcing the search to find alternative paths that do not mis-assemble.

GGAssembler segments the target gene sequence into variable fragments that encode diversity and constant ones devoid of diversity. We provide an output table that specifies the variable fragments and the sequences of the constant fragments that can be custom-synthesized or amplified from a preexisting gene.

We next introduce a streamlined wet-lab protocol (**Figure 2**) for assembling a library generated by GGAssembler(Hoch et al., n.d.). To maintain cost-effectiveness, we amplify constant fragments with primers flanked by the appropriate restriction-enzyme recognition site and all variable DNA oligos with a single 3’ primer, to allow filling the reverse strand by a polymerase. By adjusting the concentration of each DNA fragment in the assembly reaction the user can bias the resulting library towards or away from specific variants or mutations providing high operational freedom.

### Accurate assembly of vHH combinatorial libraries

We verify the accuracy of GGAssembler by constructing two complex camelid antibody (vHH) libraries, each comprising nine DNA segments. Unlike GFP, vHHs are small (<130 amino acids), allowing us to analyze the accuracy and representation bias of the library through deep sequencing.

Using the htFuncLib protein design approach that generated the GFP libraries(Weinstein et al. 2023), we designed two camelid antibody (vHH) libraries, vHH1 and vHH2, that encoded diversity in the three complementarity-determining regions (CDRs) of each antibody. Briefly, htFuncLib uses Rosetta atomistic design calculations(Leaver-Fay et al. 2011) and a machine-learning method called EpiNNet(Lipsh-Sokolik et al. 2023) to select mutations that are predicted to combine freely to generate low-energy multipoint mutation libraries. This approach circumvents experimental mutation scanning and nominates mutations for combinatorial variant libraries that are predicted to be tolerated both individually and in combination with one another. We demonstrated that such designed variant libraries are enriched in stable, well-folded, and potentially functional protein variants that exhibited vast changes in functional profile(Weinstein et al. 2023; Lipsh-Sokolik et al. 2023). Applied to the vHHs, htFuncLib results in 55,296 and 233,280 designs, in 11 and 13 positions across the three CDRs for vHH1 and vHH2, respectively. Following protein design calculations, we applied GGAssembler, including codon compression.

We transformed the libraries into bacterial cells and sequenced 12 and 14 colonies for vHH1 and vHH2, respectively. 75% and 93% of the resulting sequences were in frame and matched desired designs for vHH1 and vHH2, respectively, on par with the current state of the art(Daffern et al. 2023; Choi et al. 2022). In both errors observed in vHH1, a pair of overhangs misligated, resulting in a short product **(Supplementary Table 1)**. Retrospectively, the overhangs used to generate vHH1 were designed based on data from an early study on fidelity in GGA(Potapov, Ong, Kucera, et al. 2018); but a subsequent study(Potapov, Ong, Langhorst, et al. 2018) showed that the selected gates exhibit low ligation fidelity (60%) matching the fidelity observed in Sanger sequencing (**Figure 4A**). Using the updated fidelity data(Pryor et al. 2020; Potapov, Ong, Langhorst, et al. 2018) in GGAssembler, the overhangs in vHH1 would not have been selected.

**Figure 4.**
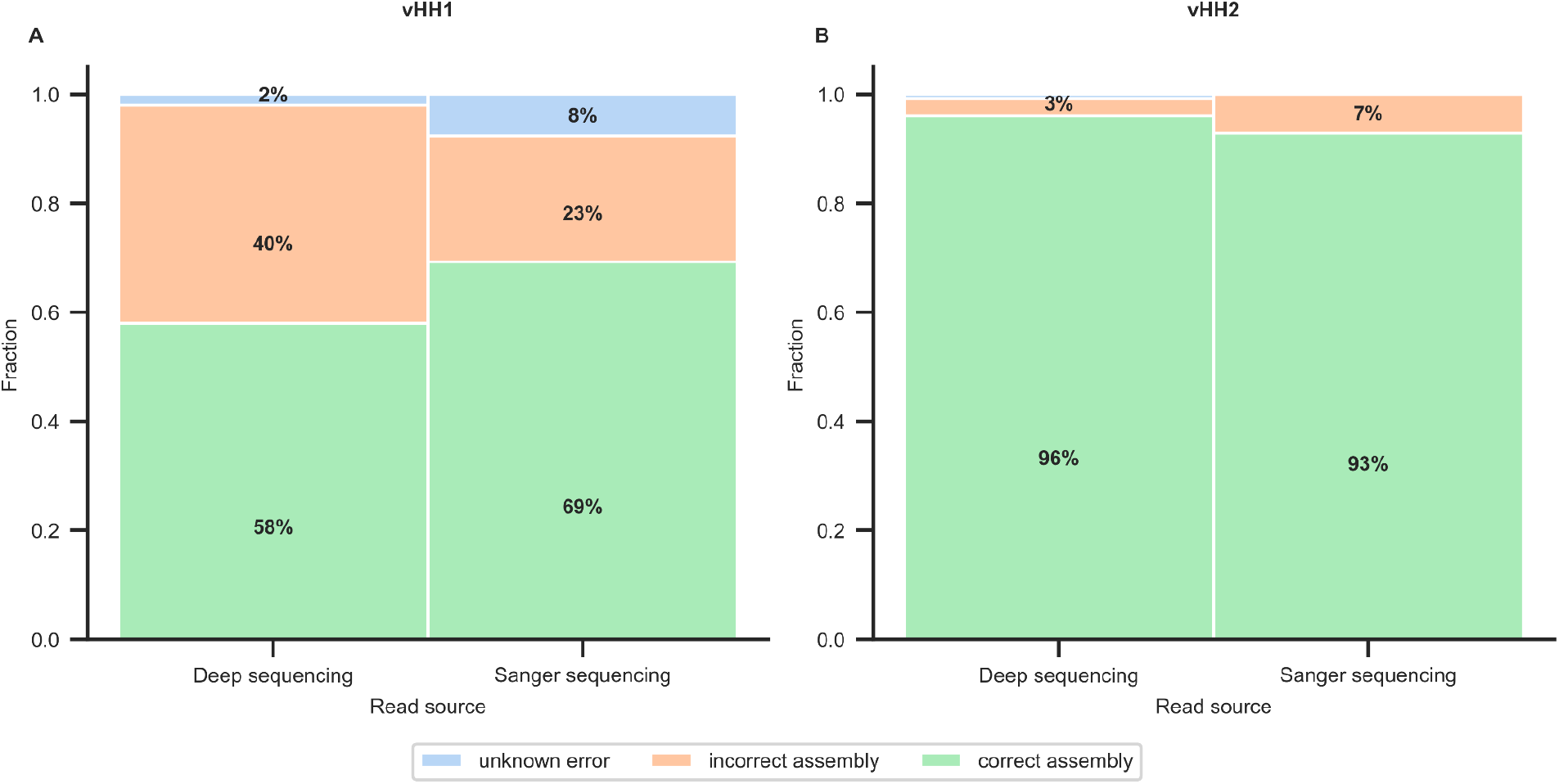
Sequencing read classification. (**A, B**) Reads were classified based on their alignment to each of the fragments. We classified reads containing both flanking constant segments based on the type of error we found “incorrect assembly”, “correct assembly” and “unknown error”. We have omitted class fractions below 1%.

Next, we subjected the assembled libraries (before transforming into bacteria so as not to introduce biological bias) to deep sequencing analysis. To assess biases in the library we started with stringent quality control, discarding any read that was not completely identical to a desired variant in any part of the variable segments (mismatches and deletions were allowed in the constant segments). 96% and 93% of designed variants were uniquely identified in vHH1 and vHH2, respectively, resulting in nearly complete coverage of the designed libraries. The distribution of variants is expected to be multimodal, and the observed distribution of variants reflects the expected mixed Poisson distribution very closely (**Figure 5A and B**). Only two variants exhibited more than a tenfold count difference in vHH1 (out of >55,000 variants) and approximately 1,300 of vHH2 (out of >230,000 variants; 0.6%), and in both libraries, the discrepancy was at most 22 fold (**Figure 5A and B, insets**).

**Figure 5.**
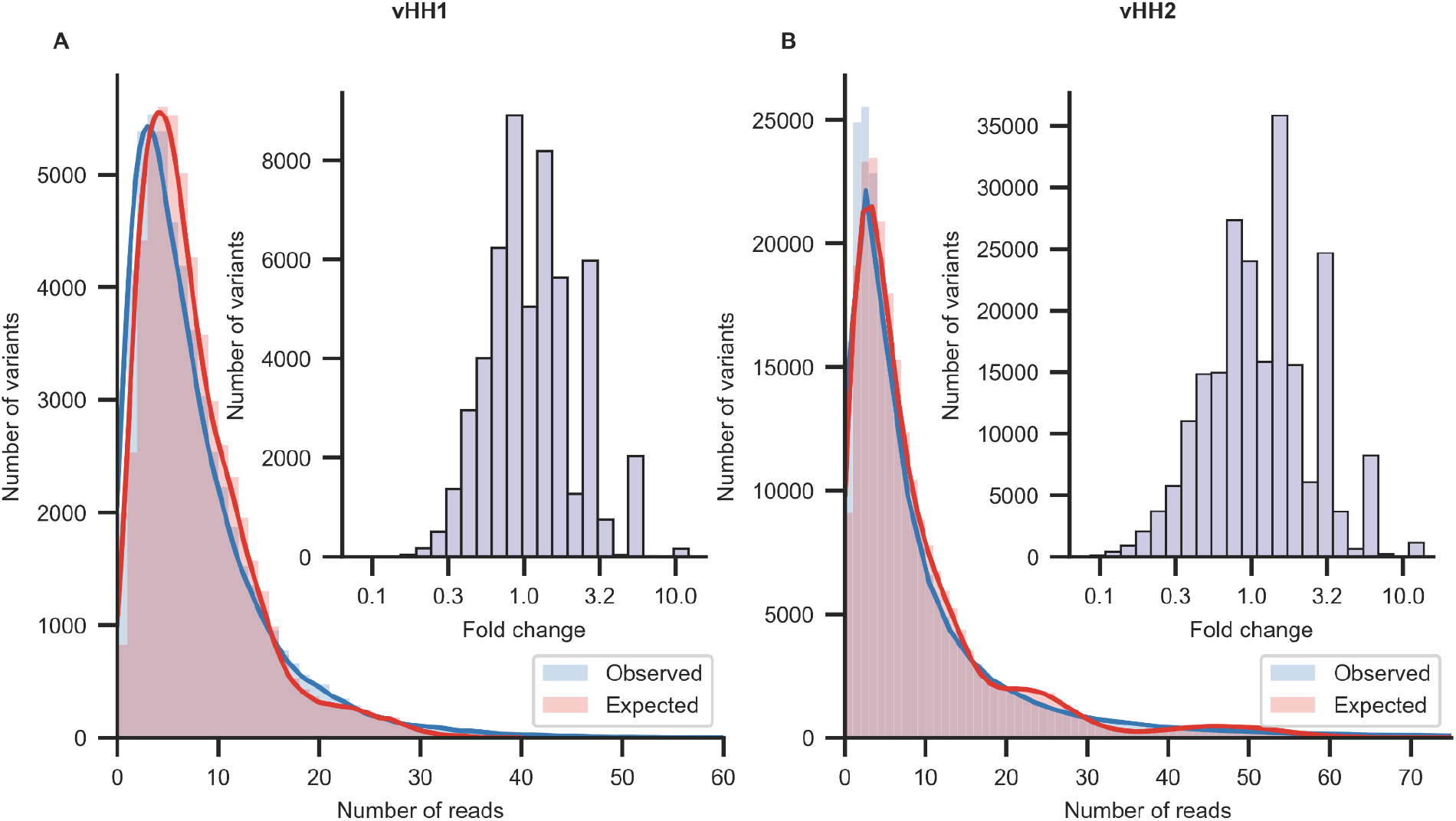
Low bias in GGAssembler libraries. Deep sequencing analysis of libraries (A) vHH1 and (B) vHH2. The observed distribution of variant reads compared to the expected mixed Poisson distribution. (insets) Log fold differences between expected and observed read counts.

We also assessed the observed frequencies of variable segments relative to the expectation. The fold-change differences ranged between 0.58-1.76 and 0.47-2.67 in vHH1 and vHH2, respectively (**Figure 6A, B**). Mutation frequencies exhibited an even narrower range, with fold-change ranging 0.67-1.26 and 0.72-1.32 in vHH1 and vHH2, respectively (**Figure 6C, D**); this range is considered to be near-uniform(Daffern et al. 2023).

**Figure 6.**
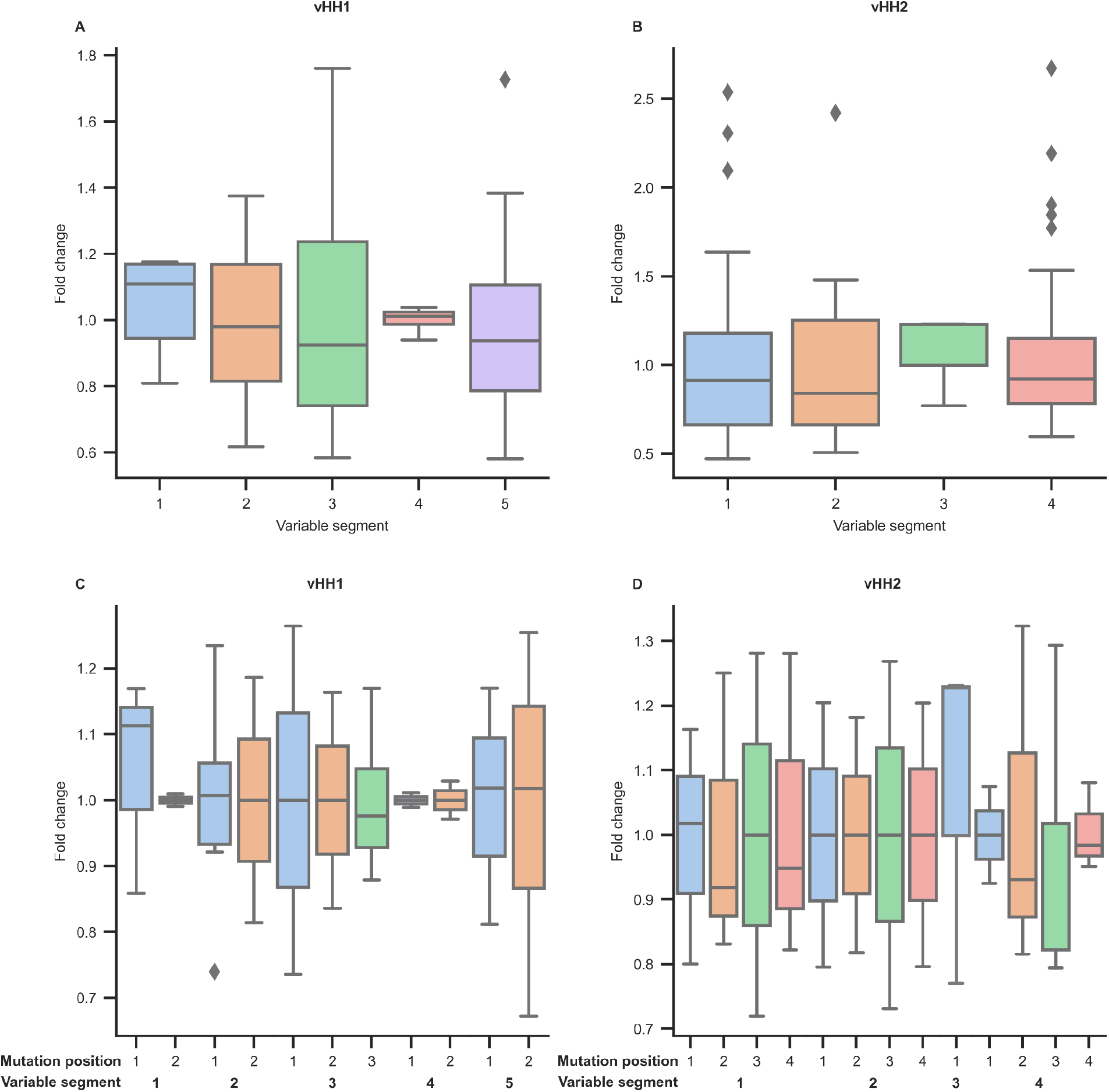
Low-bias in the representation of assembled segments and mutations. (**A, B**) Observed versus expected distribution of segments in the assembled product. The vHH1 and vHH2 libraries were encoded with 5 and 4 variable segments, respectively (for cost effective representation of all diversity in CDRH3). Comparison of the theoretical frequency of each variable segment compared to the observed frequency seen in deep sequencing. (**C, D**) Observed versus expected distribution of mutated positions. In all plots, box bounds signify the first, median and third quartiles. Whiskers represent 1.5 times the interquartile range and outliers are shown as diamonds.

Finally, we analyzed misligations to understand the reasons for failures in assembly (see Methods) (**Figure 4A, B, Supplementary Table 2**). Correct and incorrect assembly products were distributed similarly to the results of the Sanger sequencing. Incorrect assemblies (40% and 3% for vHH1 and vHH2, respectively) were all shorter than the desired assembly. Additionally, <3% of the sequencing reads were assigned as misassembled for unknown reason. The distribution of correctly and incorrectly assembled product is similar in Sanger and deep sequencing (**Figure 4A, B, Supplementary Table 2**), highlighting again the high accuracy of the method and the importance of using the more recent and reliable GGA empirical ligation data(Potapov, Ong, Langhorst, et al. 2018; Pryor et al. 2020) as implemented in vHH2. We conclude that GGAssembler produces a nearly complete representation and low bias of the designed variants and correct assembly of the vast majority of the product. Thus, the GGAssembler method and the experimental assembly protocol generate highly complex variant libraries reliably, accurately, and economically.

The total reaction costs for vHH1 and vHH2, respectively, are $392.16 and $439.08 of which $365.85 and $425.70 (93% and 97%) are DNA-associated costs, translating to 0.7₵ and 0.2₵ per variant **(Supplementary Table 3)**. Costs scale roughly logarithmically with the number of variants, with larger libraries exhibiting a substantially reduced cost per variant; for instance, the number of variants in vHH2 increased by 4.2 fold relative to vHH1 whereas the total library costs only rose by 12%. We further note that in many variant screening workflows, one may wish to generate alternative libraries that focus diversity on different regions of the protein. In such cases, parts of the assembly reaction can be reused to generate differentially biased products at no additional DNA cost.

## Discussion

New approaches to probe the mutational tolerance of proteins experimentally and computationally fuel interest in methods for efficient synthesis of combinatorial mutation libraries(Tretyachenko et al. 2020; Wrenbeck et al. 2016; Kirby et al. 2021; Daffern et al. 2023; Jacobs et al. 2015; Shimko, Fordyce, and Orenstein 2020; Püllmann et al. 2019; Yamamoto et al. 2020; Alejaldre, Pelletier, and Quaglia 2021; Sidore et al. 2020; Plesa et al. 2018; Öling et al. 2022; Lund et al. 2024). GGAssembler exploits the fact that desired diversity in active sites is typically clustered in several contiguous epitopes (*e*.*g*., antibody CDRs or enzyme active-site loops) that are separated by immutable regions. This clustering lends itself to breaking the assembly reaction into constant fragments that can be synthesized or amplified from a preexisting gene and variable regions that can be economically encoded by custom-synthesized short oligos or oligo pools. GGAssembler uses codon compression and chooses a set of empirically determined high-fidelity overhangs to minimize costs and ensure accurate assembly even in complex libraries with more than a dozen mutated positions across several noncontiguous epitopes. In the two camelid antibody examples provided here and our previous application to the GFP chromophore-binding pocket(Weinstein et al. 2023), correct assemblies were made from 9 and 20 parts, respectively, demonstrating that even large, multi-epitope active sites can be effectively generated with a simple experimental workflow that can be accomplished in a one-pot *in vitro* reaction. Critically, our variant libraries were designed strictly according to atomistic considerations without attempting to reduce the complexity of the assembly reaction like previous variant library studies(Jacobs et al. 2015; Shimko, Fordyce, and Orenstein 2020); yet the GGAssembler approach successfully produced economical, high-fidelity libraries. Thus, GGAssembler opens the way to encoding user-defined diversity in large and complex functional proteins.

While we strove for generality in developing GGAssembler, we note that it only applies to constructing combinatorial assembly libraries. With the current active research pushing the boundary of what is possible with GGA, we hope to extend GGAssembler capabilities further to allow the assembly of multiple homologous proteins in the future. Our results show the importance of inferring overhang fidelity from experiments under settings used in the assembly reaction(Pryor et al. 2020; Potapov, Ong, Kucera, et al. 2018), with notable improvement in predictive ability for library vHH2 relative to to the previously assembled vHH1 library. Future research into the fidelity of multiple restriction enzymes and ligases in the same reaction pot could enable executing complex multi-step reactions in a single-step. We envision that GGAssembler will enable efficient and economical study of the mutational tolerance of antibodies, enzymes, pathogenic antigens and other functional proteins and an effective approach to screen for new or improved functions(Lipsh-Sokolik et al. 2023; Weinstein et al. 2023).

## Methods

### Golden gate assembly optimization

The fidelity *F* of a given overhang *O* is defined as the proportion of the number of times overhang *O* ligates to its Watson-Crick pair and vice versa (*N*_*correct*_) divided by the number of times overhang *O* ligates to any overhang (*N*_*total*_).

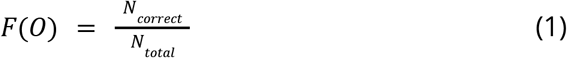

The efficiency of a given overhang *O* is relative to overhang with the most number of ligation events *O*_*max*_ and defined as the proportion of of times overhang *O* ligates to any overhang (*N*_*total*_) divided by the the number of times overhang *O*_*max*_ ligates to any overhang (*N*_*max*_).

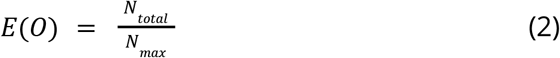

### Computational Library design

#### Codon compression

For each diversified amino acid position, we codon compressed the required diversity by generating all ambiguous codons that encode a portion of the required diversity using CodonGenie(Swainston et al. 2017) and formulating the compression as the set cover problem, where the desired amino acid diversity in each position is the set to be covered, and each ambiguous codon provided by CodonGenie is a subset of that required diversity. We have implemented Dancing Links (DLX)(Knuth 2000) to solve the exact cover problem, which results in every amino acid identity appearing exactly once.

#### DNA segmentation

We model all possible segmentations as a directed graph with each node representing a cleavage site and overhang, and edges representing DNA segments between a pair of cleavage sites. Edges are weighted by the number of nucleotides required to produce that segment to represent relative DNA costs. As stated in the results, finding economical segmentations that assemble correctly is a known NP-hard problem (Lozano and Medaglia 2013). To overcome the hardness barrier, we developed a randomized algorithm employing color-coding(Alon, Yuster, and Zwick 1995), drawing random node colorings that color nodes with the same color if empirical ligation preferences(Potapov, Ong, Kucera, et al. 2018; Pryor et al. 2020; Potapov, Ong, Langhorst, et al. 2018) indicate the overhangs would ligate. Next, for each random coloring drawn, we run a rainbow version of the Dijkstra shortest-path algorithm(Dijkstra 1959), which considers extending a rainbow path only if the added node color does not already appear in the path. The result is a DNA segmentation and set of overhangs that ensure high-fidelity assembly. Selecting rainbow shortest-path solutions guarantees the assembly of an economical solution that excludes undesired products.

Our proposed solution is a randomized algorithm for which we will show that bounding the probability of an error, either not finding a path or finding the wrong path, requires an exponential dependence only on the number of fragments. In practice, we found that running the algorithm for e^#fragments^ iterations ensures finding the lowest cost solution with high probability, if such a solution exists. In many cases, the difference in oligo cost between the lowest-cost solution found and other solutions is negligible (can be on the order of tens of bps). With such a slight difference in cost, we prefer solutions with fewer fragments or higher fidelity. We have also observed that even running for 10^2^-10^3^ iterations, the solutions are close in cost to the best solution found, resulting in a run-time of several minutes from start to finish. We will now prove the run-time complexity of the algorithm.

#### Algorithm proof

Rainbow Dijkstra (CD) algorithm:

##### CD(G, c, s, t)

1. Initialize a priority queue (min-heap) to store vertices based on their distance from the source *s*. Each element in the priority queue will be a triplet of (distance, vertex, color set), where the color set represents the set of colors used to reach that vertex.
2. Start from vertex *s*. Initialize its distance as 0 and its color set as {0}.
3. Repeat until the priority queue is empty:
  a. Pop the vertex *v* that is closest to the priority queue.
  b. For each neighbor *w* of *v*:
    i. If *w* shares a color with the *v* color set, skip this neighbor.
    ii. Otherwise, calculate the new distance to *w* as the distance to *v* plus the weight of the edge from *v* to *w*.
    iii. If this new distance is shorter than the current distance to *w*:
      1. Update the distance to *w*.
      2. Update the color set of *w* as the union of the color set of *v* and the color of *w*.
      3. Add *w* to the priority queue.
4. Once *t* is reached, the shortest rainbow path from *s* to *t* is found.

We will begin by analyzing the runtime complexity of a rainbow version of the Dijkstra shortest-path algorithm.

###### Lemma 1

Let graph *G* = (*V, E*) be a directed or undirected graph with vertex set *V* and edge set *E*, and let *c: V* → {1, ⋯, *k*} be a coloring of its vertices with *k* colors, *w: E* → *R*_≥0_ an edge weight function and *s, t* ∈ *V*. A rainbow shortest path of length at most *k* from *s* to *t* in G, if one exists, can be found in *O*(|*E*||*V*|) worst-case time.

**Proof:** Using a priority queue (min-heap) implementation, the Dijkstra algorithm has a time complexity of *O*((∣*V*∣ + ∣*E*∣)*log*∣*V*∣). For each vertex, we may need to examine all its neighbors to find the shortest path [*O*(∣*E*∣) operations]. For each neighbor, we may need to check the color set [*O*(|*V*|) time]. Thus, the overall time complexity for finding the shortest vertex rainbow path from *s* to *t* is: *O*((∣*V*∣ + ∣*E*∣)*log*∣*V*∣ + ∣*E*∣.∣*V*∣) = *O*(|*E*|.|*V*|).

Rainbow shortest path (CSP) algorithm:

##### CSP(G, k)

1. To find a rainbow shortest path of length *k* in *G* that covers the entire WT DNA, we add two vertices, a source vertex *s* and a target vertex *t* colored by 0 and connect *s* to the vertices in *G* and the vertices in *G* to *t* such that the new edges obey the restrictions set by the user.
2. Loop:
  a. Draw a random vertex coloring *c: V* → {1, …, *k*}.
  b. Run a rainbow version of Dijkstra’s shortest-path algorithm and record the result.

Since CSP is a randomized algorithm we are interested in bounding the probability of an error, either not finding a shortest path or that the solution found is not optimal.

###### Lemma 2

Let graph *G* = (*V, E*) be a directed or undirected graph with vertex set *V* and edge set *E*. Bounding the probability of an error at ε requires

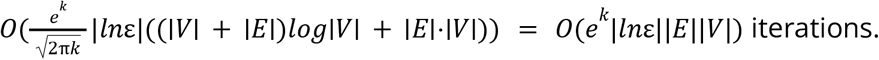

**Proof:** Let *P* be the shortest vertex rainbow path of length *k*. The probability of selecting a random coloring *c: V* → {1, …, *k*} for which *P* is a rainbow path is 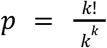 and so for a single iteration of the loop the probability of an error is 1 − *p*. To bound the probability of CSP to result in an error by at most ε we require *t*(ε) iterations. We will now solve for *t*(ε).

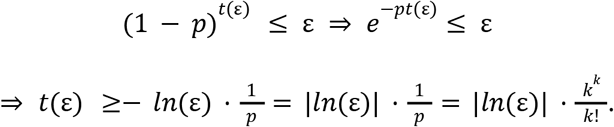

Applying Stirling’s approximation we get:

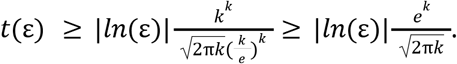

Having shown that the number of iterations required to bound the probability of error by ε is 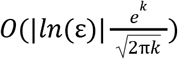 and because each iteration of the rainbow version of the Dijkstra shortest-path algorithm requires *O*((∣*V*∣ + ∣*E*∣)*log*∣*V*∣ + ∣*E*∣.∣*V*∣) time, we conclude that the run time in order to bound the probability of error by ε is

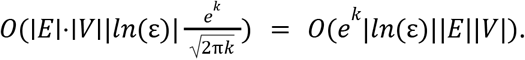

### Experimental Library Assembly

The full protocol is described in ref. (Hoch et al., n.d.).

The primers used for the reactions are described in Supplementary Tables 4 and 5.

### Library Cloning

The assembled library after Golden Gate was cloned into pNACP plasmid(Uchański et al. 2019) (kindly provided by Jan Steyaert, Vrije Universiteit Brussels) using homologous recombination in yeast(Gietz and Schiestl 2007). 100μl yeast cells after recombination were plated on SDCAA plates. Colonies were collected with 1.5mL sterile DDW and extracted using Zymoprep Yeast Plasmid Miniprep II (Zymo Research, CAT #D2004).

#### Colony PCR and Sanger sequencing

The cloned libraries after recombination to pNACP were transformed into E. Cloni 10G cells (Lucigen). Next, colony PCR and Sanger sequencing were performed using standard protocols.

### Deep Sequencing

Amplicon libraries were prepared as previously described(Blecher-Gonen et al. 2013). The libraries after GGA were sequenced using a paired-end V3 600 cycles Illumina kit (Illumina MS-102-3003) on an Illumina Miseq. We preprocessed Fastq sequences using the BBTools software suite(Bushnell, Rood, and Singer 2017), and pairwise aligned the results to constant and variable DNA segments using Parasail(Daily 2016). We further analyzed alignment results using Python(McKinney 2010; The pandas development team 2023; Harris et al. 2020; Van Der Walt, Colbert, and Varoquaux 2011; Cock et al. 2009; Granger and Pérez 2021), by assigning each read bp its corresponding segments. This allows us to quickly identify correctly assembled sequences and assign each incorrectly assembled sequence the reasons it was assembled as such. We then extracted variable segments and filtered out sequences that did not exactly match variable DNA segments for the bias analysis.

## Acknowledgements

We thank David Peleg for helpful discussions and comments on the algorithm and Olga Khersonsky and Ariel Tennenhouse for critical reading. Figures 1, 2 and 3 were created with BioRender.com. The research was supported by the Volkswagen Foundation (94747), the Israel Science Foundation (1844), the European Research Council through a Consolidator Award (815379), the Dr. Barry Sherman Institute for Medicinal Chemistry, and a donation in memory of Sam Switzer.

## Notes

### Competing Interest Statement

The authors have declared no competing interest.

### Summary of Updates

Added a section on Algorithm proof; Revised Figure 1 and 2 caption; Added a colab notebook for easier use.

